# Parallelized Droplet Vitrification Enables Single-Run Vitrification of the Whole Rat Liver Hepatocyte Yield

**DOI:** 10.1101/2024.07.14.603471

**Authors:** M. S. Taggart, A. Tchir, L. Van Dieren, H. Chen, M. Hassan, C. Taveras, AG. Lellouch, M. Toner, R. D. Sandlin, K. Uygun

## Abstract

Drug discovery pipelines rely on the availability of isolated primary hepatocytes for investigating potential hepatotoxicity prior to clinical application. These hepatocytes are typically isolated from livers rejected for transplantation and subsequently cryopreserved for later usage. The gold-standard cryopreservation technique, slow-freezing, is a labor-intensive process, with significant post-storage viability loss. In this work, we introduce parallelized droplet vitrification, a technique for high-volumetric, rapid vitrification of suspended cells. We show the utility of this technique through the single-run vitrification of the whole-rate liver hepatocyte yield, resulting in a 1600% increase in single-batch vitrification and a 500% increase in droplet generation rate compared to previous droplet vitrification approaches. Additionally, we showed that these implementations maintained improved post-preservation outcomes in primary rat hepatocytes.

## Introduction

The current standard in drug discovery pipelines is using high-throughout, bulk screening utilizing commercially available, cryopreserved hepatocytes for studying drug metabolism and toxicity to identify hepatotoxicity early in drug development^1–4^. Primary human hepatocytes are typically isolated from transplant-rejected, non-viable livers since transplantable livers are typically allocated to a recipient. These organs are initially thought to be potentially transplantable and determined to be non-viable following procurement. Hepatocytes are then isolated through a perfusion-based collagenase extraction, after which they are rapidly stored to prevent deterioration while maintaining functionality, enabling later usage. The current pool of liver graft donors is largely homogenous, with disparities existing in donation by sex, race, and age, resulting in a stark shortage of available donor livers for hepatocyte isolation in underrepresented groups^5^. Additionally, since livers tend to be procured from donors with potential transplantability, livers from individuals with various pathologies such as metabolic disease are not typically procured or isolated^6^. These factors, when considered together, outline the current challenges in drug discovery, where most drugs are screened on primary hepatocytes derived from relatively healthy donors, and the results are therefore generalizable only for healthy individuals, representing a narrow demographic. A key limiting factor for the usage of suboptimal livers is a lack of cryopreservation techniques that retain high viability and yields. The current standard for hepatocellular storage, slow-freezing, results in suboptimal cell yields and poor viabilities, leading to inefficiencies in the usage of donor cells, and exacerbating the already stark shortage of diverse donor cells^7^.

The current gold standard for primary hepatocyte cryopreservation is by slow-freeze process, involving a pre-storage incubation phase with cryoprotective agents (CPAs) to reduce intracellular-ice-formation (IIF)^8^. Briefly, isolated hepatocytes are suspended at 5 x 10^6^ – 1 x 10^7^ cells/mL in a storage solution; typically, University of Wisconsin (UW) with added albumin and a monosaccharide for improved oncotic/osmotic stability, although newer solutions with improved efficacy, have been reported^9^. The cells are kept on ice at 4C and after suspension, are incubated with 10% dimethyl sulfoxide (DMSO) for 10-15 minutes, during which a shrink-swell cycle occurs due to an osmotic shift whereby intracellular water is replaced by DMSO^10^. Following this incubation phase, the cells are cooled to -80C at a controlled rate to minimize the nucleation of damaging ice. Despite wide-spread adoption and standardization of the slow-freeze technique, the stresses undertaken during freezing and thawing result in considerable cellular damage^11^. To avoid such stresses, an alternative cryopreservation approach is vitrification, where cells are cooled quickly through the glass transition that allows ice nucleation to be bypassed^12^. One such application, bulk droplet vitrification, is a technique that employs this strategy in the storage of primary rat hepatocytes via the introduction of cell-laden “droplets” directly onto a liquid nitrogen (LN_2_) surface. By employing a rapid-mixing stage before droplet formation, CPA concentrations are achieved sufficiently high to enable vitrification, while cellular toxicity is avoided^13^. A pilot trial using this technique showed great success at the scale of 25 million cells, yet limitations with the volumetric capacity remained, preventing its applicability in the storage of human hepatocyte yields, which can exceed 100 billion.

Working toward the goal of single-run vitrification of human liver hepatocytes, whereby the entire, bulk hepatocyte yield is vitrified in a single batch, we developed parallelized droplet vitrification, a scalable technique for large-volume vitrification. Through the introduction of a parallel-flow, splitting phase before CPA introduction, the single-run throughput was increased to encompass the entire rat liver hepatocyte yield of approximately 250 million cells/run (which is improved over the previous iteration using 25 million hepatocytes), while the volumetric flow rate was increased to 10 mL/min, an improvement from 2 mL/min. The splitting design is inherently modular, enabling the vitrification of greater volumes through the addition of further parallel splits. Additionally, we introduced a multi-tube collection device to enable the storage of multiple aliquots of vitrified cells, without the necessity of post-vitrification manipulation which decreases the risk of spontaneous ice formation.

## Methods

### Experimental Design

Parallelized droplet vitrification (PDV) was compared to both slow-freeze cryopreservation and standard small-scale bulk droplet vitrification (ss-BDV). Additionally, the efficiency of each technique was compared to freshly plated cells (within 3 hours of isolation). Immediate post-isolation fresh cell viability and 24-hour culture viability were evaluated from ≥5 isolations. For slow-freeze and ss-BDV, immediate post-preservation yield and viability were determined from ≥5 individual aliquots, while 24-hour culture viability was determined from ≥3 platings for each group. Both immediate post-preservation yield and viability, as well as 24-hour culture viability for PDV, were determined from 1 individual aliquot, plated in eight wells. Vitrified droplet characterization was performed on 4 individual aliquots.

### Primary Rat Hepatocyte Isolation

All animals used in this study were approved under protocol #2011N000111 by the Institutional Animal Care and Use Committee (IACUC) of Massachusetts General Hospital. The animals were housed socially in a temperature and humidity-controlled room and provided unrestricted access to food and water. Primary rat hepatocytes were isolated from adult, female Lewis rats (10-12 weeks old, 150-250g) (Charles River Laboratories, Wilmington, MA USA) as previously described^14^. Briefly, rats are anesthetized under isoflurane and the abdomen is opened. Following dissection of liver connective tissue, the portal vein is cannulated and 200 mL of oxygenated 2.12 uM EDTA in KRB is perfused to completion at 17 mL/min to, followed by 200 mL of 0.12 mg/mL of collagenase. The hepatocytes are then extracted in KRB using forceps, filtered through a 250 um filter, followed by 90 um, and spun down at 25G for 5 mins. The cells are then resuspended and spun down in percoll at 50G for 10 mins. Finally, the cells are resuspended in DMEM for plating and cryopreservation.

### Droplet Vitrification

ss-BDV is performed as previously described^13^. Following isolation, 40 million hepatocytes were spun down at 25G for 5 mins and resuspended on ice in University of Wisconsin solution (UW) (Bridge to Life, Duluth, GA USA) containing 2.4 mg/mL bovine serum albumin (BSA) (Sigma-Aldrich, St. Louis, MO USA). A dimethyl sulfoxide (DMSO) (Sigma-Aldrich)/ethylene glycol (EG) (Sigma-Aldrich) mixture is then added sequentially at 7.5% v/v and 15% v/v, with a 3-minute incubation phase for each concentration, resulting in a final cell concentration of 10M/mL. At the end of 3 minutes, 10 uL of the cell suspension is taken and cell concentration and viability are determined. The cells are then loaded into a 3 mL syringe. A second 3-mL syringe is loaded with UW containing 2 mg/mL BSA, 800mM sucrose (Sigma-Aldrich), and 65% v/v EG/DMSO. The syringes are then placed in a custom adapter and placed in a syringe pump (Pumpsystems, Kernersville, NC USA). The syringe tips are connected via .078” x .125” tubing (Radnoti, Covina, CA USA), leading to a mixing needle (Grainger, Lake Forest, IL, USA) ending in a 24G needle (Becton Dickinson, Franklin Lakes, NJ, USA). The pump is angled vertically so the needle faces directly down at a dewar (Sigma-Aldrich) filled with LN_2_. Inside the dewar is a 3D-printed funnel connected directly to a 50 mL conical. The pump is run at 2 mL/min for 3 min, resulting in a total volumetric flow of 5 mL, with 2.5 mL from each syringe, resulting in a cellular suspension with final CPA concentration of 40% DMSO/EG and 400 mM sucrose. The conical cap is punctured with a 24G needle, and parafilm is wrapped around the top to trap vapor nitrogen, resulting in maintenance of the cryogenic temperature. Parallelized droplet vitrification follows the same workflow, however, several changes are made to the vitrification apparatus to enable greater volumetric flow rates **(Fig. 1a)**. A custom-made branching device is implemented at the outflow **(Fig. 1b)**. To allow for increased flow-rate, each syringe is connected to 16G tubing, which branches into 4 parallel channels of .078” x .125” tubing to allow for 4 parallel flow channels. Each separate flow channel ends in a mixing nozzle connected to 24G needles, enabling 4 distinct droplet generation sites. A 3D-printed 4-way funnel is placed in the dewar **(Fig. 1c,d)**. The funnel is connected directly to separate 50 mL conicals, resulting in the single-run production of 4 vitrified droplet aliquots **(Fig. 1e)**. The same EG/DMSO, BSA, and sucrose concentrations are used for the cell-suspension and high-CPA syringe as optimized from the ss-BDV protocol, however, 100 million cells are suspended in a total 11 mL and the solutions are suspended in 10 mL syringes. The syringes are connected to the custom branching device, and the pump is run at 6 mL/min, resulting in consistent droplet production **(Fig. 1f)**.

**Figure 1:**
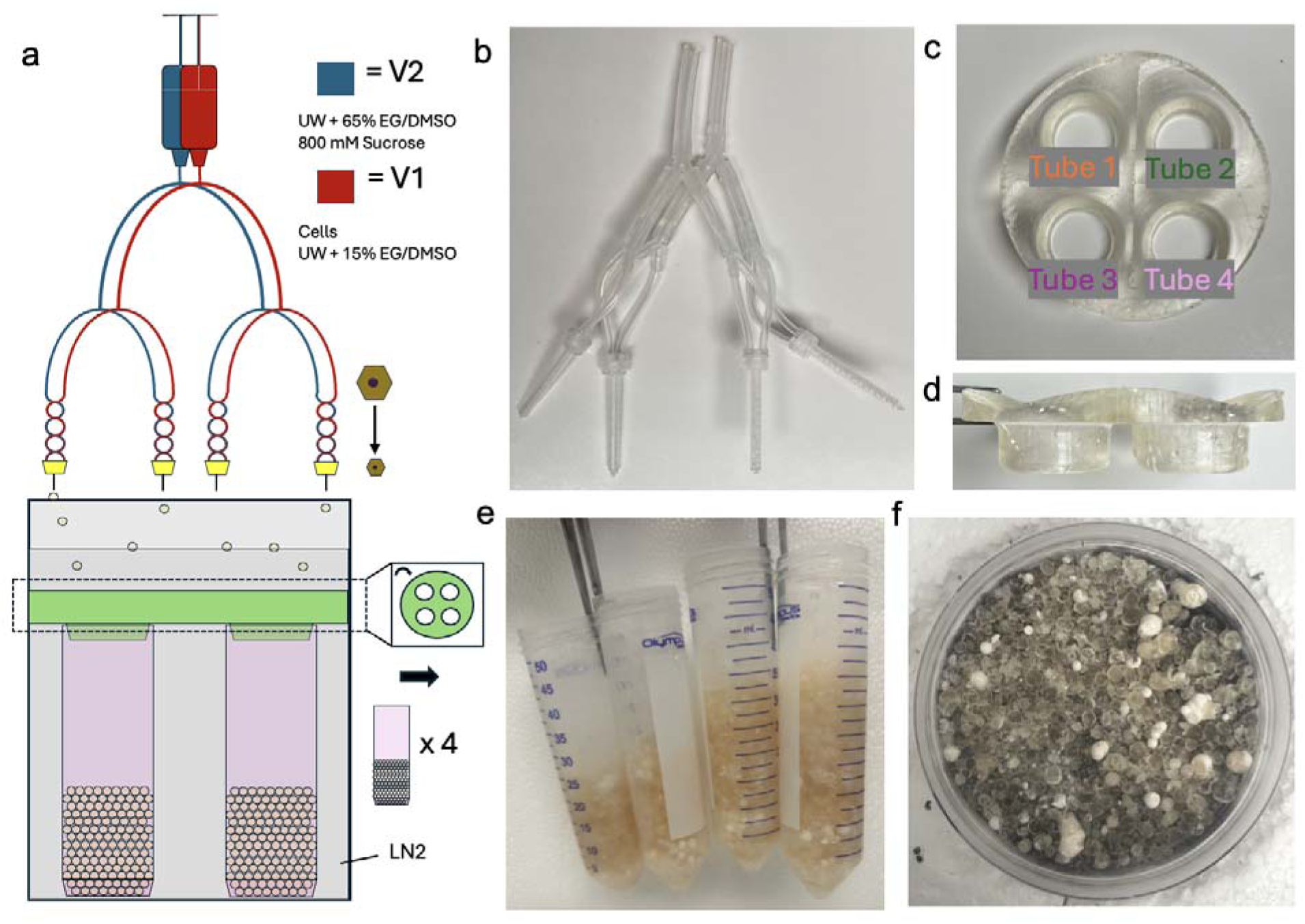
Design of a Scalable, Parallelized Droplet Vitrification System. Process outline displaying the workflow of a scalable droplet vitrification device **(a)**. Two syringes containing 10mL of cell solution (V1) and vitrification solution (V2) are run on a syringe pump at 6mL/min through tubing of descending size, after which the separate solutions enter a helical mixing needle and exit through 24G needles **(b)**. The droplets fall onto a liquid nitrogen surface, on which they vitrify and drop into a 3D-printed droplet sorter **(c, d)**. resulting in equal droplet distribution between four 50mL conicals **(e)**. A sample of droplets with a high vitrification percentage, demonstrated by degree of transparency; frozen droplets are opaque and milky white **(f)**.

Since PDV and ss-BDV result in aliquots of similar volume, the same rewarming process is used for both. 100 mL of DMEM supplemented with 500mM sucrose is warmed to 37C in a water bath. A conical containing vitrified droplets is removed from storage and placed in a LN_2_ container. Rapidly, the droplets are poured into the bottom of a 250 mL beaker and immediately warmed by DMEM poured on top. The suspension is simultaneously stirred until all droplets are rewarmed. Cells are then rapidly spun down in 2 separate 50 mL tubes at 50G for 10 min, after which 37.5 mL is aspirated and the cells are resuspended via gentle rocking. To gently rehydrate the cells and prevent osmotic shock, 12.5 mL and 25 mL DMEM are added sequentially with a 3-minute acclimation period between. Finally, the cells are spun down at 25G for 5 minutes and resuspended in 4 mL DMEM.

### Vitrified Droplet Characterization

To characterize the droplets, a custom black background was made from a divot in Styrofoam, enabling the droplets to be kept at -196C while under a microscope **(Fig. S1a)**. The back of a petri dish is blackened, and a scale is etched into the surface **(Fig. S1b)**. The divot is filled with LN_2_ and the petri dish is floated on top, with a thin layer of LN_2_ inside. The vitrified droplets are then poured from the 50 mL conical into the petri dish and imaged using a digital camera. To process the images, they were opened in ImageJ (NIH, Bethesda, MD USA), an internal scale was set based on the etching, and the diameter of individual droplets was manually determined, assuming sphericity **(Fig. S1c)**. To determine whether droplets were frozen, they were visually inspected and sorted, with frozen droplets being opaque white, and vitrified droplets being transparent with a brown tint due to the presence of hepatocytes **(Fig. S1d)**.

### Slow-Freeze Cryopreservation

Isolated cells are spun down at 25G for 5 minutes and resuspended in 4C UW supplemented with 2 mg/mL BSA and 100 mM D-glucose (Sigma-Aldrich). 10% v/v DMSO is added to the suspension resulting in a final cell concentration of 10 million cells / 1.5 mL and incubated for 20 minutes. After 20 minutes, a 10 uL aliquot is taken and the concentration and viability of cells is determined. Following the DMSO incubation, the cells are aliquoted to cryotubes at 1.5 mL/tube and moved to a controlled rate freezer and frozen using the following scheme: starting at 4C, cool 1C/min to 0C and hold for 8 mins, after which, cool 2C/min to -8C, then 35C/min to - 28, 2.5C/min to -33C, warm 2.5C/min to -28C, cool 1C/min to -60C, finally cool 10C/min to - 100C. After cooling, the frozen tubes are rapidly moved to deep cryogenic storage at -196C. To rewarm, the cryotubes are removed from storage and thawed into a 37C water bath. To prevent cracking from thermal stress, the tube is gradually submerged until only a small piece of ice remains within the cell mix, at which point it is brought to a biosafety hood and kept on ice. The cell suspension is added to a 50 mL conical containing 6 mL isotonic percoll and 14 mL DMEM, inverted, and spun down at 50G for 5 mins. The pellet is then resuspended in 2 mL DMEM and kept on ice.

### Cell Culture

Hepatocytes were cultured on Collagen 1 treated 24-well plates (Corning, Corning, NY, USA). Cells were suspended in DMEM at 500k cells/mL or 700k cells/mL, for fresh and cryopreserved cells respectively and 500 uL of media was added to each well. After addition, the plates were shaken aggressively to allow for the cells to spread throughout the well. The plates were then moved to the incubator and the cells were allowed to adhere for 1 hour or 1.5 hours for fresh and cryopreserved cells respectively. After adherence, the cells were washed with warm DMEM, and finally, incubated overnight in C+H.

### Cell Concentration and Viability Determination

A 10 uL aliquot of cell suspension is taken and mixed at a 1:1 ratio with Trypan Blue solution (Gibco, Waltham, MA USA). 10 uL of the mixture is taken and added to a hemocytometer, after which the live and dead cells are counted. This process is done immediately prior to cryopreservation, and immediately after. To determine the yield, the ratio of the total number of cells observed following cryopreservation and prior to cryopreservation is determined according to the following equation:

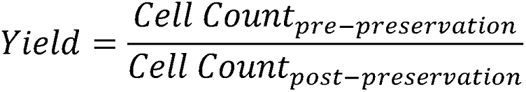

To determine viability, the ratio of the number of live cells to total cells is determined according to the following equation:

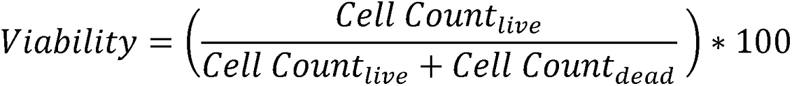

### Plated Cell Viability Determination

Cells are stained with the Live/Dead mammalian cell viability kit (Invitrogen, Waltham, MA USA) as well as NucBlue nuclear probes (Invitrogen) 24 hours after plating, according to the manufacturer’s instructions. Briefly, the cells are washed with Dulbecco’s Phosphate Buffered Saline (DPBS) (VWR, Radnor, PA USA) followed by light-sensitive incubation at 21C for 30 minutes in 200 uL DPBS containing 2 uM calcein AM, 4 uM Ethidium homodimer-1, and 2 drops/mL NucBlue. Following incubation, the cells are washed with and resuspended in DPBS. The cells are then imaged on an EVOS microscope and quantified using FIJI **(Fig. S2a)**.

To determine the plated viability, the following equation was used:

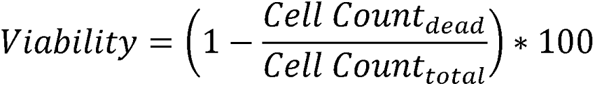

### Mathematical Simulations

The volumetric changes over time were simulated as done previously^13^. Briefly, we used the Kedem-Katchalsky (K−K) formalism^15^:

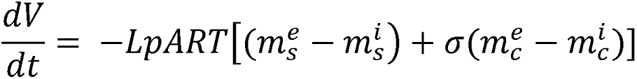

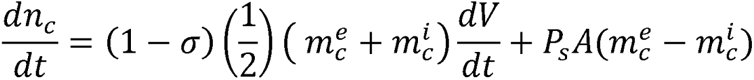

Where V is the cell volume, A is the surface area, and n_c_ denotes the intracellular CPA content. L_p_ stands for hydraulic conductivity, P_s_ indicates the membrane permeability to CPA, and σ is the reflection coefficient. R represents the gas constant, and T is the absolute temperature. m refers to the molality, with superscripts “i” and “e” specifying intracellular and extracellular environments, respectively, and subscripts “s” and “c” indicating nonpermeating salts and permeating CPA, respectively. The coupled ordinary differential equations described above were solved using Python in Jupyter Notebook

The critical warming and cooling rates were calculated as described previously^16^. Briefly, the multispecies critical cooling rate was calculated based on the following formula:

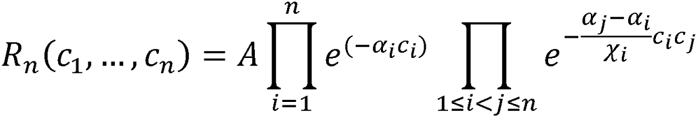

Where A and α are constants and the pre-exponential and exponential factor, respectively. C is the weight fraction concentration and R is the critical cooling or warming rate. Pre-exponential and exponential factors for DMSO, EG, and sucrose were retrieved from the literature^17^.

### In Silico Simulations

For modeling the cooling rates and times according to droplet size, we used COMSOL Multiphysics® (version 5.5, Comsol AB, Stockholm, Sweden) as done previously^18^. The initial temperatures of the droplet and LN_2_ were set to 4 °C and −196 °C, respectively. A convective heat flux was established as the boundary condition between the droplet and the cold LN_2_, with a natural convective heat transfer coefficient of 100 W m^−1^ K^−1^. To model the convective warming temperature profile, the initial temperature of the droplet was set to −196 °C, and a convective heat flux was set as the boundary condition between the droplet and the rewarming medium (37 °C), with a forced convective heat transfer coefficient of 500 W m^−1^ K^−1^. Cooling and warming rates were calculated between -20 and -140 degrees Celsius.

### Statistical Analysis

Droplet size data within individual tubes was compared using a paired t-test. We assessed cross-tube comparisons using one-way ANOVA with multiple comparisons. Post-preservation data was analyzed using one-way ANOVA with multiple comparisons. The droplet-size, viability relationship was analyzed by a simple linear regression. Plated cell stain quantification was also compared using one-way ANOVA with multiple comparisons. All one-way ANOVA comparisons were performed with Tukey’s multiple comparisons test, with single pooled variance. Stars denote significance: *0.01 < p < 0.05; **0.001 < p < 0.01; ***0.0001 < p < 0.0001; ****0.0001 < p. Analysis, as well as graphing was done on GraphPad Prism version 10.0.3 (GraphPad Software, Boston, MA USA).

## Results

### Characterization of Resultant Vitrified Droplets

To characterize the parallelized droplet vitrification process, droplets were counted and their size was determined following vitrification. Additionally, to determine the consistency of droplet collection, variations between collection tubes was determined. Each collection tube had an average of 127 ± 5.5 droplets, of which, 112 ± 4.5 were vitrified and 17 ± 6.4 were frozen, accounting for 86.9% ± 4.5 and 13.1% ± 4.5 of the droplet population respectively, showing significantly more vitrified droplets compared to frozen droplets (p = 0.0009) **(Fig. 2a)**. The size distribution of the droplets was consistent between collection tubes, with an average size difference of 43% ± 0.04 between vitrified and between vitrified and frozen droplets **(Fig. 2b)**. Overall, the average diameter of vitrified droplets was 2.66 mm ± 0.83 while frozen droplets had an average diameter of 4.13 mm ± 0.84, revealing a significant difference in diameter between frozen and vitrified droplets (p < 0.0001) **(Fig. 2c**). Additionally, the proportion of droplets vitrified beyond 4 mm in diameter made up only 6.5% of all vitrified droplets, whereas 56.0% of frozen droplets were greater than 4 mm in diameter **(Fig. 2d)**.

**Figure 2:**
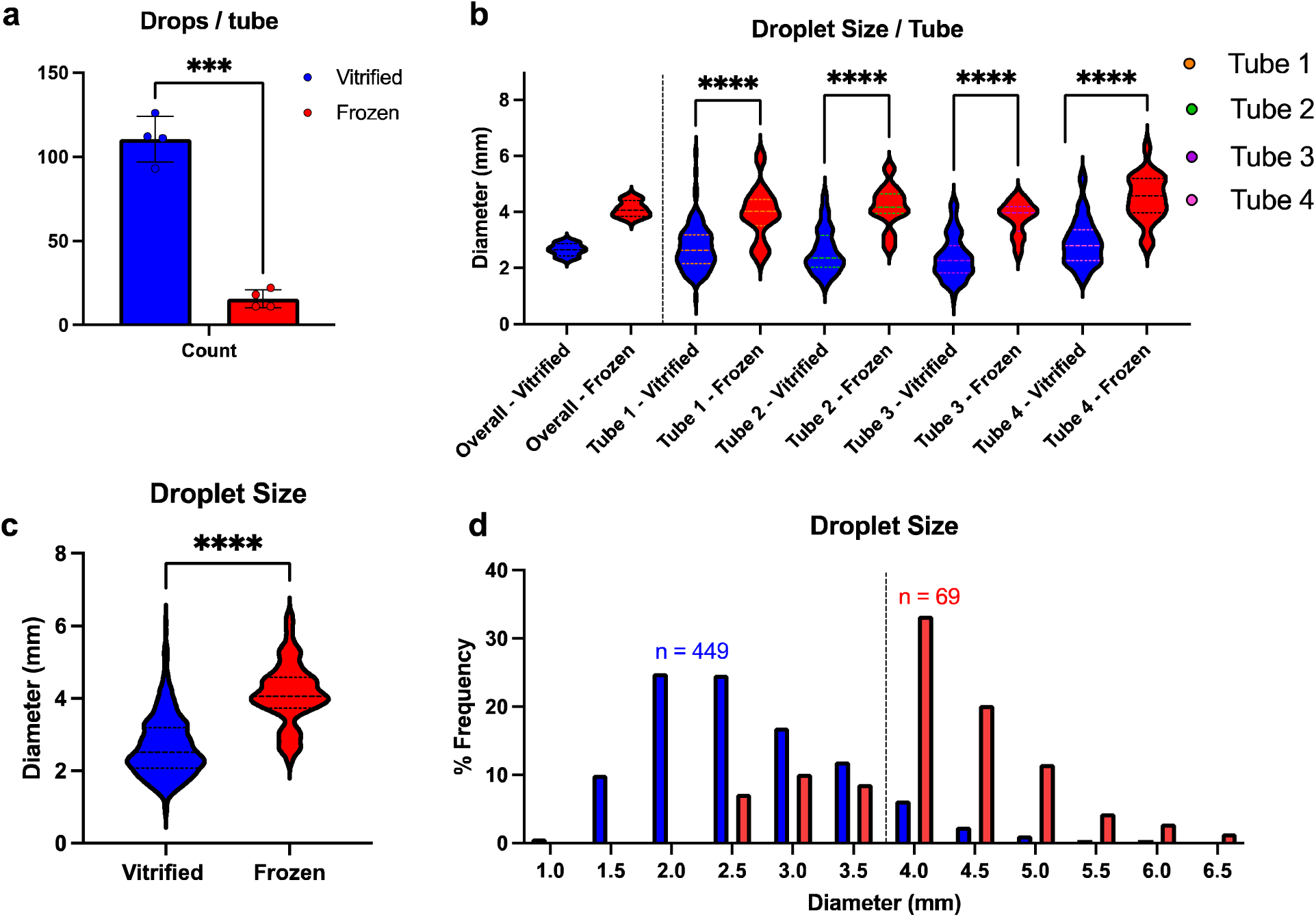
Vitrified Droplets show a Distinct Size Threshold. Characterization of droplets obtained through parallelized droplet vitrification of 100 million cells. Tubes showed an average vitrified count of 110.5 ± 13.5 droplets, constituting 87.8% ± 3.7 of the sample population (n = 4) **(a)**. Droplet size showed a consistent trend across all four tubes, with an average size difference of 43% ± 4.4 between vitrified and frozen droplets **(b)**. When comparing vitrified droplets (2.7mm ± 0.8) to frozen droplets (4.1 ± 0.8), a significant size difference is observed (p < 0.0001) **(c)**. Droplet size appears to correlate with successful vitrification, resulting in a threshold of 4 mm, beyond which few droplets vitrify **(d)**. Results are displayed as mean ± standard deviation.

### Modeling of the Droplet Vitrification Process

To gain insight into cellular and droplet dynamics throughout the droplet vitrification process, the process was simulated using a mathematical model. It was found that during the two CPA incubation steps, cells recover to their full initial volume, whereas, upon the final mixing step, they rapidly shrink in response to the osmotic gradient, resulting in nonequilibrium vitrification **(Fig. 3a)**. The critical cooling and warming rates were determined and compared to the simulated rates of 2 mm and 4 mm droplets at varying CPA concentrations. It was determined that within the final CPA concentration range, 2 mm droplets fell within the ice-free, vitrification zone, whereas 4 mm droplets fell inside the crystallization zone **(Fig. 3b)**. It was also determined that the warming rate of both droplets fell well outside the vitrification zone, explaining bulk ice recrystallization upon rewarming **(Fig. 3c)**. To gain deeper insight into the role of droplet size on vitrification, we simulated the cooling and warming rate of a range of droplet sizes, finding an inverse relationship between droplet size and heat transfer rate; darker droplet represents those produced by the bulk droplet vitrification process **(Fig. 3d)**. The cooling of several droplet sizes was simulated, and it was determined that both 2 mm and 3 mm droplets show approximate homogenous cooling, taking approximately 7 and 10 seconds respectively to reach -140°C, whereas heterogeneous cooling was observed in 4 mm droplets, with the periphery and inside taking approximately 12.5 and 14 seconds respectively to cool **(Fig 3e)**.

**Figure 3:**
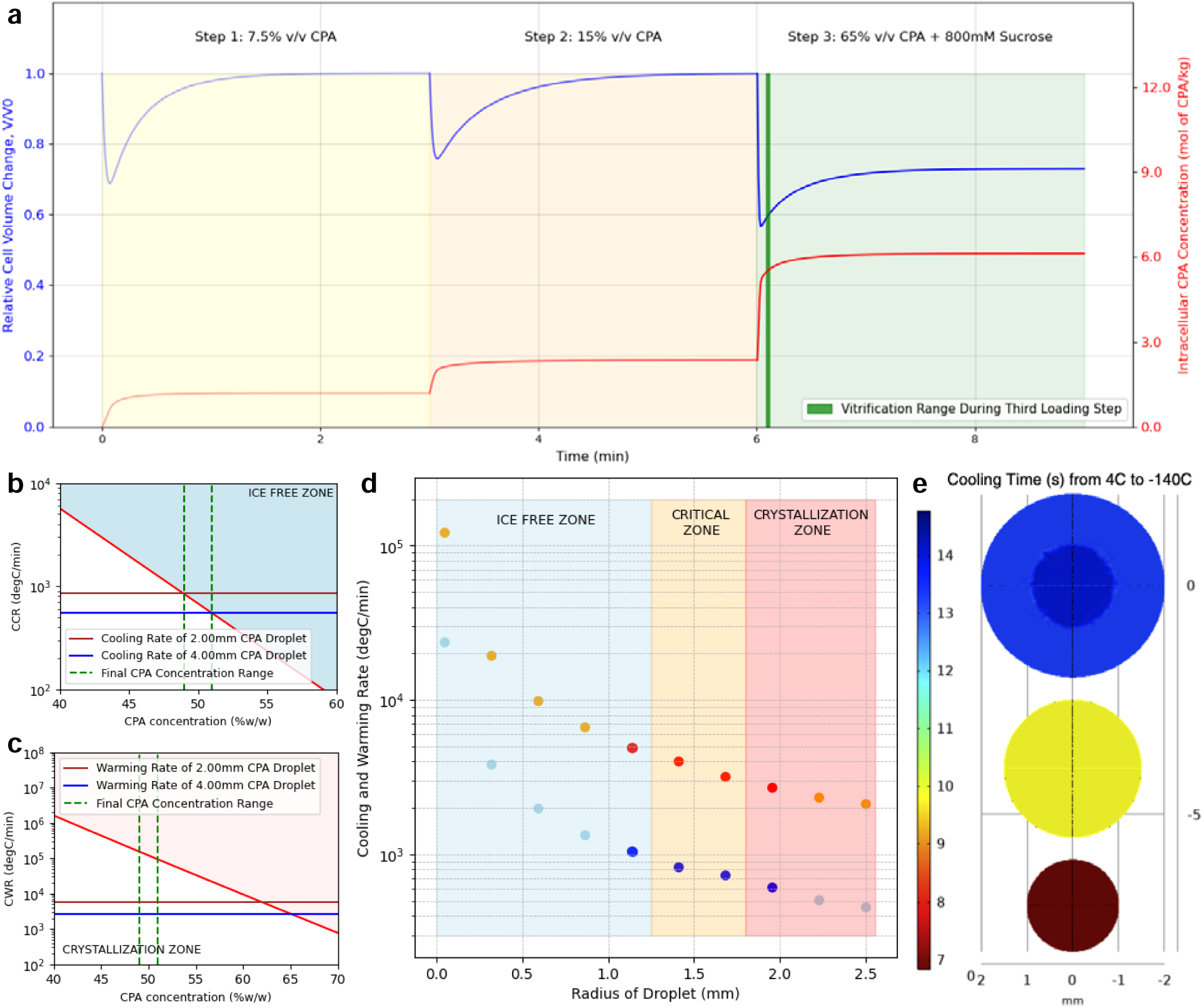
Post-hoc mathematical and in silico analysis of droplet vitrification. A mathematical model was employed to gain a better understanding of the vitrification dynamics at the droplet-LN_2_ interface. The incubation steps show size recovery of individual hepatocytes prior to the next step; however, the vitrification phase shows non-equilibrium size change and CPA uptake (**a**). The critical cooling rate for 2 mm droplets falls within the non-crystallization zone, whereas 4 mm droplets do not, explaining the size disparity between frozen and vitrified droplets (**b**). The critical warming rate confirms bulk recrystallization during the rewarming phase for both 2mm and 4mm droplets, as they fall outside the non-crystallization range (**c**). An inverse relationship between size and cooling and rewarming rate was described; darker droplets represent those observed through experimentation (**d**). The radial relationship between time to reach -140C from 4C shows heterogeneity in 4mm droplets, while 2mm droplets show approximate homogeneity (**e**).

### Immediate Post-Thaw Characteristics of Scaled Droplet Vitrification

To determine the impact of rewarming on the cells, post-thaw characteristics were determined. The yield was slightly elevated for large-scale bulk droplet vitrification (PDV, 84.2% ± 33.4) compared to 64.1 ± 7.1 (p = 0.1639) for small-scale BDV (ss-BDV) and 63.3% ± 6.8 (p = 0.1326) for slow-freeze **(Fig. 4a)**. The immediate post-thaw viability was determined for each group and compared to the viability of 92.8% ± 4.3 for fresh cells following the isolation resulting in 88.1% ± 1.8 for slow-freeze (p = 0.6135), 68.5% ± 9.2 for ss-BDV (p < 0.0001), and 69.6% ± 8.0 for PDV (p = 0.0006); additionally, a significant difference was observed between slow-freeze and PDV (p = 0.0031) **(Fig. 4b)**. The immediate viability was determined for droplets individually rewarmed, resulting in a median viability of 62.8%, with an IQR of 12.5% **(Fig. 4c)**. To determine the homogeneity of cells within each droplet, the number of total cells recovered from each rewarmed droplet was counted, resulting in a median cell count of 402,000 cells/droplet with an IQR of 346,000 cells/drop, with several outliers resulting from droplets merged during vitrification **(Fig. 4d,e)**. The relationship between immediate post-thaw viability and total number of cells recovered in each droplet had low correlation (r^2^ = 0.05436) **(Fig. 4f)**.

**Figure 4:**
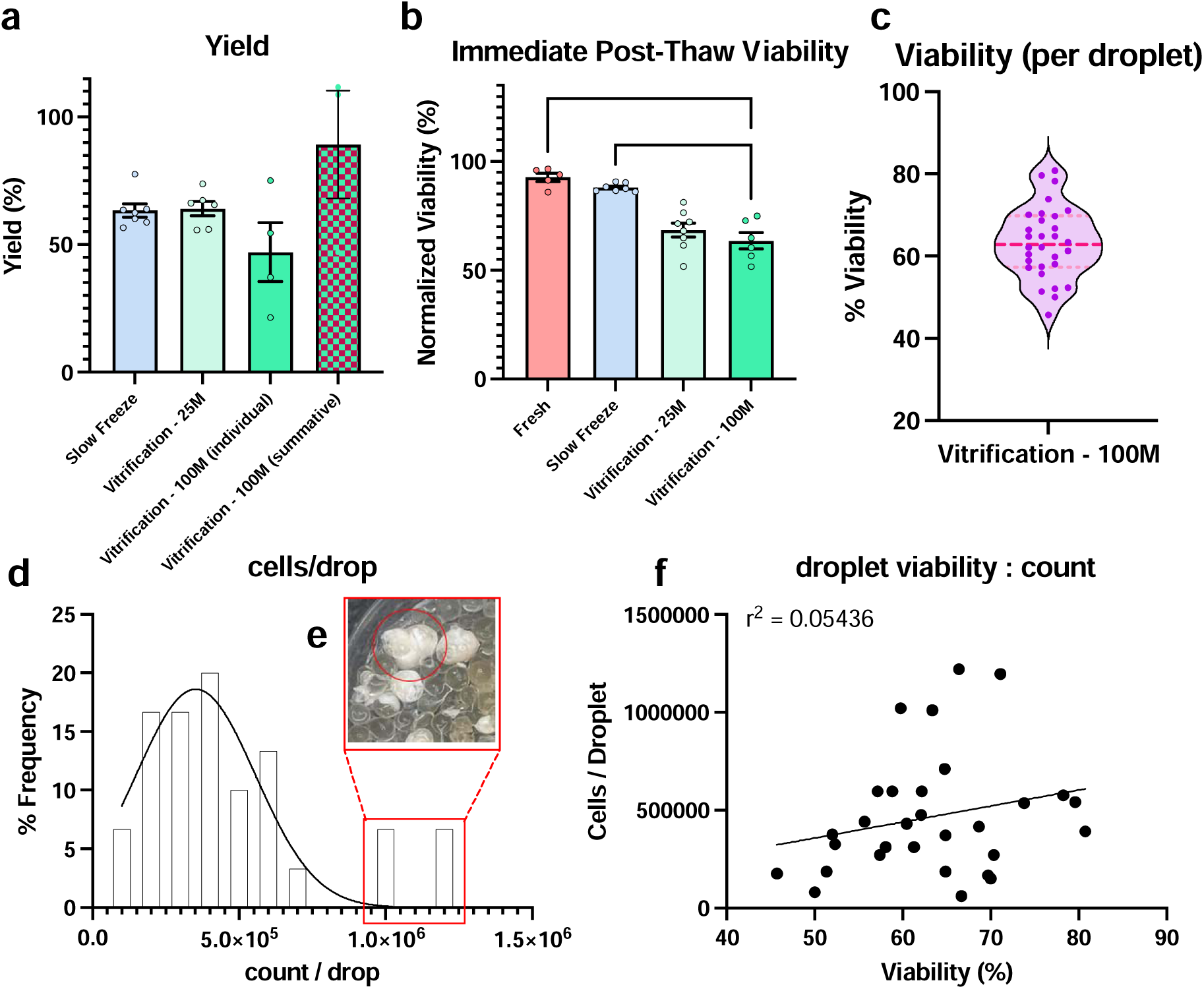
Immediate Post-Thaw Characteristics of Scaled Droplet Vitrification. Cells cryopreserved using slow freezing (gold standard), standard bulk droplet vitrification (BDV, 25M cells), and parallelized droplet vitrification (PDV, 100M cells) were thawed and compared. Immediately following thawing, cells were counted, and the normalized live yield was determined. PDV resulted in a comparable yield to both slow-freeze and BDV. Data is presented for both individual tubes (individual) and for each run (summative) **(a)**. Additionally, immediate normalized viability showed no difference between approaches; each data point is from a separate replicate/separately processed batch of cells and looks at the total population viability **(b)**. From a single PDV run, individual droplets (n = 30) were thawed, and immediate cell viability was determined, revealing a median of 62.8%, IQR 12.5 **(c)**. Individual droplets had a median cell count of 402,000 cells/drop, with an interquartile range of 346,000 **(d)**. Droplets with high cell counts (>100,000), may be accounted for by droplets that merged during vitrification **(e)**. Viability did not correlate with droplet cell count (r^2 = 0.05436) **(f)**.

### Parallelized Vitrification Retains Hepatocyte Viability

Following rewarming, cells were plated overnight on 24-well plates to determine the post-thaw attachment efficiency and compared to freshly plated cells. The average total number of attached cells per view for fresh cells was 1385 ± 160, showing a significant reduction when compared to 599 ± 103 for slow-freeze, 712 ± 104 for ss-BDV, and 953 ± 305 for PDV (p < 0.0001 comparing fresh to each preservation method). A significant improvement in total cells per view was observed for PDV when compared to slow-freeze (p = 0.0002) **(Fig. 5a)**. The average number of dead attached cells per view for fresh cells was 34.9 ± 29.9, showing a significant reduction when compared to 130.1 ± 34.4 for slow-freeze (p = 0.0008), and 182.6 ± 97.1 for ss-BDV (p < 0.0001), while no difference was observed when compared to 37.9 ± 30.1 for PDV (p = 0.9994). A significant reduction in total dead cells per view was observed between slow-freeze and PDV (p = 0.0060) **(Fig. 5b)**. The viability of the plated cells for each group was determined by finding the ratio of dead cells to total cells per view. There was no significant difference in plated/cultured viability between fresh hepatocytes 95.8% ± 3.8 and PDV 95.5% ± 4.1 (p > 0.9999). Fresh cell viability was significantly elevated compared to 76.2% ± 7.8 for slow-freeze (p < 0.0001)), and 74.0% ± 12.7 for ss-BDV (p < 0.0001). Additionally, a significant increase in plated viability was observed between PDV and slow-freeze (p = 0.0001) as well as PDV and ss-BDV (p < 0.0001) **(Fig. 5c)**. Looking directly at hepatocyte morphology, fresh hepatocytes showed complete coverage of the plate, with typical, hexagonal morphology, further supported by green fluorescence from live staining. Strong bile canaliculi formed between cells in regions of high confluency in each group. Intercellular connections were greatly exaggerated in PDV compared to slow-freeze due to greater density, indicated by bright white fluorescence between cells. Live staining covered a greater region in PDV compared to slow-freeze, further supporting the fact that a greater number of healthy cells, and a lower number of dead cells attached in PDV compared to slow-freeze.

**Figure 5:**
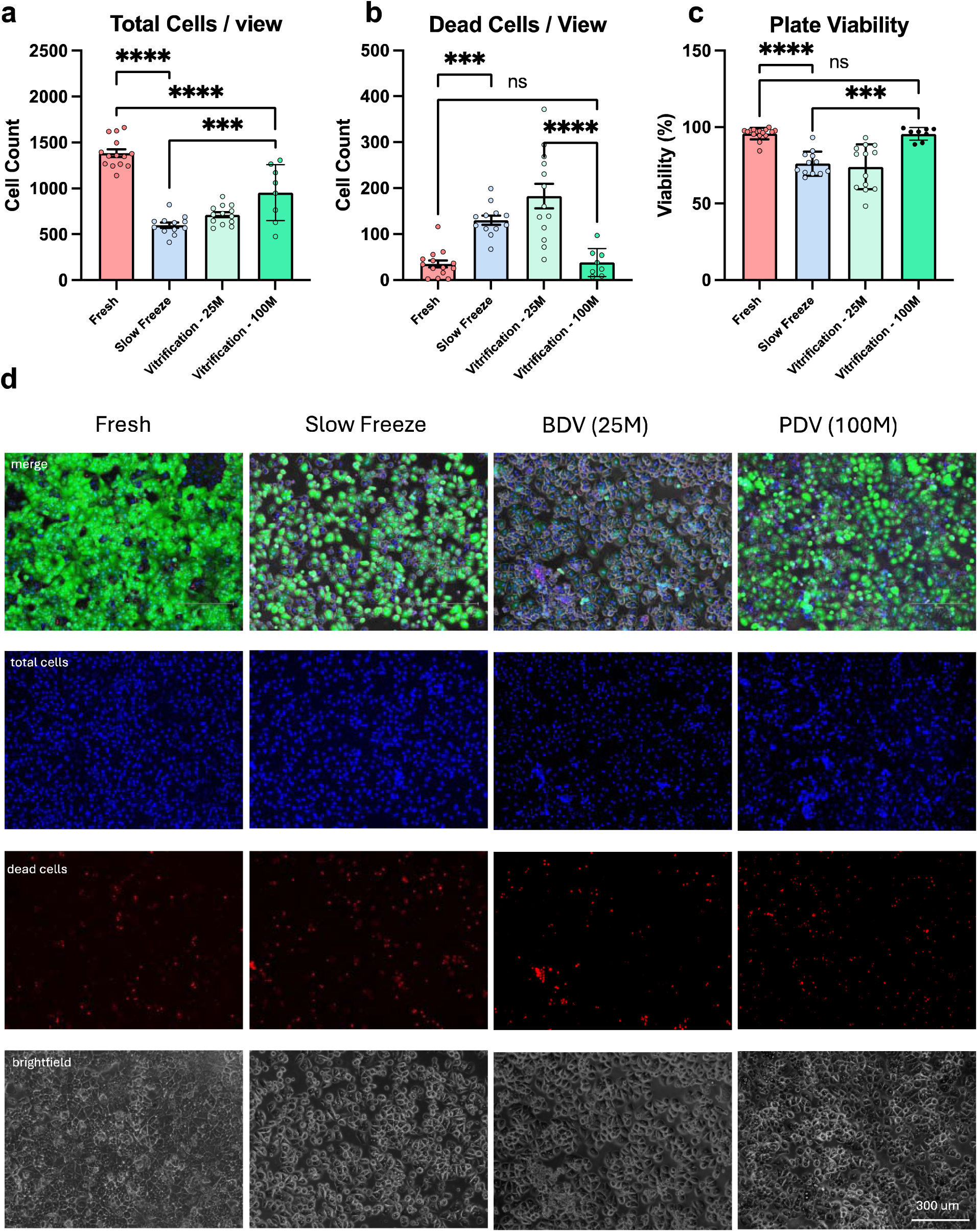
Parallelized Droplet Vitrification Results in Retained Hepatocyte Viability. Following rewarming, cells were plated on 24-well plates at 500k live cells/mL. After 24 hours of attachment, cells evaluated. PDV resulted in improved viability (95.8% ± 3.8) compared to both slow freeze (76.2% ± 7.8) and ss-BDV (74.0% ± 14.7) (p < 0.0001 for each) **(a)**. The total number of attached cells/view was significantly greater following PDV (952 ± 305) compared to slow freeze (599 ± 103) and ss-BDV (712 ± 103) (p = 0.0002, 0.0146 respectively), indicating improved attachment efficiency **(b)**. Of attached cells, PDV had fewer dead cells (38 ± 31) compared to slow freeze (130 ± 34) and ss-BDV (183 ± 97), (p = 0.0060, < 0.0001 respectively) **(c)**. Brightfield imaging reveals fewer empty patches and greater cellular density **(d)**. Error bars represent standard deviation.

### Figure 6: Single-Run, Parallelized Droplet Vitrification of the Whole Rat Liver Yield

Following validation of PDV, the whole rat liver hepatocyte yield of 250M cells was vitrified (WL PDV) **(Fig. 6a)**. The vitrified droplets were characterized **(Fig. S3)**, finding no difference in vitrification rate for 100M at 86.9% ± 4.5 and WL PDV at 86.5 ± 2.5 (p = 0.7569) **(Fig. 6b)**. The average diameter of vitrified droplets was 2.65 mm ± 0.23 for 100M and 2.80 mm ± 0.03 for WL, while the average diameter of frozen droplets was 4.11 mm ± 0.3 for 100M and 4.58 mm ± 0.42 for WL; no difference was observed for either (p = 0.7666 for vitrified, p = 0.0923 for frozen) **(Fig. 6c)**. Following thawing, the live yield was determined, with an average yield of 12.3% ± 3.9 based on total vitrified count, or 49.0% ± 15.6 based on the assumption each tube should contain 25% of the total yield (WL theory). WL theory live yield was not significantly reduced compared to 63.3 ± 6.8 for slow freeze (p = 0.3533), or 47.0% ± 23.1 100M PDV (p = 0.9965) **(Fig. 6d)**. Post-thaw viability for WL PDV was 72.8% ± 4.4 and 87.6% ± 5.3 following a percoll spin, resulting in no difference compared to fresh at 92.8% ± 4.2 or slow freeze at 88.1% ± 1.8 (p = 0.3752 and p = 0.9524 respectively) **(Fig. 6e)**. Following plating, WL total cells/view was 408.7 ± 48.9, a significant reduction compared to slow-freeze at 599 ± 103 (p = 0.0092) **(Fig. 6f)**. WL dead cells/view was 27.6 ± 21.3, significantly reduced compared to slow-freeze at 130.1 ± 34.4 (p < 0.0001) **(Fig. 6g)**. Plated viability of WL was 93.4% ± 4.7, showing no difference compared to fresh at 95.8% ± 3.8, and a significant improvement over slow-freeze at 76.2% ± 7.8 (p < 0.0001 and p = 0.5648 respectively) **(Fig. 6h)**. Cellular imaging shows sparse plating following WL PDV, however attached cells show regular hepatocyte morphology, indicating retained viability despite decreased adherence **(Fig. 6i)**.

**Figure 6:**
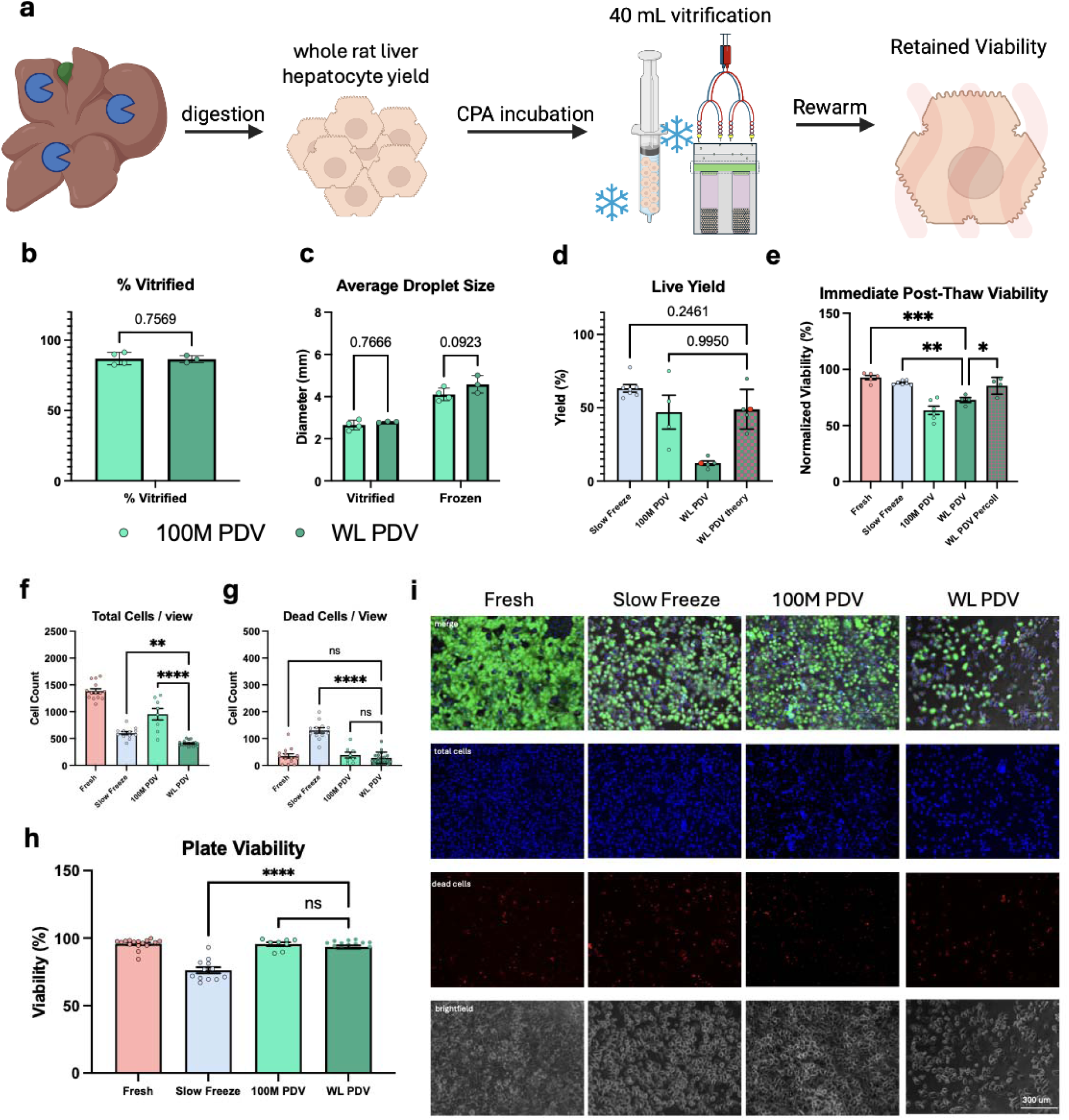
Parallelized Droplet Vitrification Enables Single Run Vitrification of the Whole Rat-Liver Hepatocyte Yield. Approaching the goal of single-run human liver hepatocyte yield, PDV was investigated for its efficacy in vitrifying the whole hepatocyte yield of the rat liver. Following isolation, the whole rat liver hepatocyte yield was CPA loaded and ran through the PDV protocol, after which, viability was determined (**A**). No difference in vitrification rate or droplet was observed (b,c). No difference in live yield was observed in 400M cells compared to 100M or slow-freeze; red data points represent the total yield for the run (**D**). Post-thaw viability was comparable to 100M PDV and, following an additional percoll spin, to slow freeze (**E**). Plated cell viability was comparable to fresh and 100M BDV (**f, g**), however a reduction in plated density was observed (**h**). Brightfield imaging reveals healthy, although sparse, cells following plating (**i**).

## Discussion

An insufficient supply of donor livers is a major bottleneck in the procurement of primary human hepatocytes for clinically translatable, *in vitro* drug discovery. Historical injustices have contributed to a wary and reluctant attitude towards organ donation resulting in further limitations in the procurement of hepatocytes from underrepresented groups^19^. This confounding effect results in disproportionate homogeneity of genetic ancestry in the stock of hepatocytes available for scientific studies, which especially limits the widespread applicability and efficacy of therapeutics to all populations^20^. To address this limitation, isolated cells must be cryopreserved in an efficacious manner to recover as many cells as possible from the limited number of livers available. The current gold standard cryopreservation technique for primary hepatocytes, slow-freezing, results in poor yields and viabilities, resulting in limited downstream usage of donor cells^13^. To address this issue, we developed a scalable, high-throughput system for bulk-droplet vitrification, resulting in an approximately 10-fold increase in single-run yield, as well as a 500-fold increase in the rate of droplet production. We characterized the droplets produced through the bulk-droplet verification technique, identifying and describing size dependence on droplet vitrification, and outlining a clear path for future developments.

To advance BDV toward clinical applications, we aimed to improve the single-run throughput through the introduction of parallel flow splitting, resulting in multiple sites of droplet production, or PDV. Following isolation, primary rat hepatocytes were suspended and underwent a 2-phase CPA incubation in which an EG/DMSO mix was introduced to the cells stepwise, enabling pre-vitrification osmotic dehydration. Dehydration of cells prior to vitrification is ideal as it minimizes intracellular water content and thereby increases the intracellular concentration of previously equilibrated CPAs, which is conducive to glass formation as opposed to ice nucleation^21^. Following pre-incubation with CPAs, the cell solution is run in parallel with a high-concentration CPA cocktail and branches out into multiple outflow streams. Parallel flow decreases the flow rate and thus shear stress experienced by the cells while allowing a higher overall droplet production rate than a single stream. Shear stress has been shown to be a detriment to hepatocyte health, resulting in both morphological and functional changes^22^. Studies have determined that beyond a critical shear domain, cells are lysed, resulting in death^23^. To enable increased overall flow rates, while minimizing shear-stress-induced damage, parallel flow was utilized to reduce the stress at each branch. Following a brief mixing step of cells with CPA cocktail to achieve desired concentration, the droplets fall directly onto LN_2_. One drawback of this setup is that the droplets may undergo a brief period of reverse Leidenfrost effect (RLE), during which they float on top of the LN_2_ due to the formation of a nitrogen air cushion formed by rapid evaporation resulting from the dramatic temperature difference between the droplets and LN_2_ which potentially leads to ice formation from a slower cooling rate^24, 25^. After sinking, the droplets are sorted by a custom-made storage device, enabling pre-storage aliquoting without the need for manual manipulation of the droplets. Future iterations of this technique may employ an advanced sorting device that aims to limit droplet size; a device with a strict, 3.5 mm radius cut-off would remove approximately 80% of all frozen droplets in this study.

We characterized the resulting droplets and found a droplet size-dependent partition between frozen and vitrified droplets. We then employed a mathematical model to show that the nature of this difference is most likely due to size-dependent cooling rates, with droplets above 4 mm not cooling fast enough to vitrify with our CPA solution. This effect is potentially confounded by the RLE. Droplets, when placed on top of LN_2_, show a strong relationship between droplet diameter and hovering time^25, 26^. Heat transfer during the hover period is rate limited due to the interfacial surface area of the droplet and the LN_2_. Due to this, droplets with a smaller radius will undergo a shorter RLE, resulting in greater heat transfer, enabling vitrification, while larger droplets undergo a prolonged RLE, resulting in heat transfer rates insufficient for vitrification. Further limiting vitrification, we observed the merging of droplets upon collision during the RLE phase, resulting in further increased diameter.

While we were able to reach vitrification rates of approximately 90%, a major limiting factor of cell viability with our current approach is the rewarming phase. Convective rewarming results in diameter-dependent heat transfer, with larger diameters resulting in decreased rates of warming in the droplet core^18^. Our modeling showed that at our CPA concentration, the droplets would need to be smaller than 1.25 mm in diameter on average to avoid recrystallization. To avoid this issue, many droplet vitrification approaches rely on low droplet volume to prevent devitrification during rewarming^27, 28^. The loss in post-thaw viability can be partially explained by the recrystallization as it is a large contributor to cell death during the post-cryopreservation thaw phase^29^. As the system volume increased, recrystallization was observed to increase as well, potentially explaining why 400M PDV resulted in lower yield and plate adherence. Recently developed methods for thawing have demonstrated rewarm rates orders of magnitude greater than convective heating^18, 30^. To prevent recrystallization and improve post-thaw viability, future studies should aim to integrate novel rewarming strategies into the PDV process, bypassing the poor rewarm rates of convective warming.

An important challenge faced when working with hepatocytes is their propensity to sediment in suspension^31^. Since hepatocytes are in a vertical syringe on the pump, we expect them to sediment towards the nozzle, resulting in unequal cell distribution between droplets. By rewarming droplets individually and finding the total cell count in each, we found a wide range of cell counts in each droplet. To determine if this effect impacted cell viability, we ran a regression on cell viability against cell count for each drop and found no relationship. This finding is beneficial to our design as it implies that cell distribution homogeneity is not required for successful vitrification. Potential disruption to this finding may be found as syringe volumes are increased, resulting in longer suspension time and increased sedimentation. In this scenario, gentle mixing technologies may be integrated to allow low-shear maintenance of solution homogeneity^32^.

A major limitation to slow-freeze cryopreservation of hepatocytes is poor post-thaw viability, resulting in decreased attachment and functionality^33^. To improve the usage of cryopreserved hepatocytes in clinically translatable work, improved post-thaw viability is a necessity. When comparing hepatocytes stored using PDV to slow-freeze, we found that PDV cells had improved viability, not significantly different from freshly plated cells. Post-thaw hepatocyte attachment is a key metric to cryopreservation success, as standard protocols result in greatly diminished plated density, downstream of reduced adhesion molecule expression^34^. PDV of 100M cells resulted in significantly enhanced attachment, with a 59% increase in cellular density compared to slow-freeze; however, a 45% reduction in density was observed compared to freshly plated cells, implying there is room for improvement in our protocol. Additionally, density reduced as the process was scaled, potentially indicating that as greater volumes are used, increased cellular injury occurs. Increased density results in greater cellular health, as evidenced by improved bile canaliculi formation, a key metric of retained hepatocellular functionality^35^. Poor attachment has been shown to be reversed in hepatic progenitor cells by the addition of hyaluronan, a stem-cell native matrix glycosaminoglycan to the cryopreservation solution, resulting in increased post-thaw adhesion molecule expression^36^. Future iterations of BDV should investigate the addition of hepatocellular matrix proteins in the vitrification solution.

While this study showed promise for the applications of PDV to primary hepatocytes, the modular nature of this technique may allow for direct translation to other cell types which have been difficult to preserve. Namely, immune cells, including NK cells, show a loss of efficacy following cryopreservation, limiting their usage as cellular therapeutics^37^. Future studies on PDV should aim to expand the suite of compatible cell types, to improve post-thaw functionality.

## Conclusion

In conclusion, we developed a scalable platform for single-run vitrification of the whole rat liver hepatocyte yield, resulting in a 10-fold increase in single-run throughput, and a 5-fold increase in vitrification rate. Our high-volume approach resulted in no difference in vitrification rate or droplet size compared to the standard technique. Following thawing, vitrified cells showed improvement in viability and adherence when plated, compared to slow-freeze. Future directions should aim to decrease the average droplet size while increasing consistency to improve the vitrification frequency and heat transfer rate. The modular, low-cost nature of this technique, holds promise for future translation and should be investigated for its efficacy in other cell lines, such as NK cells.

## Data Availability

The datasets generated during and/or analyzed during the current study are available from the corresponding author upon reasonable request.

## Author Contributions

Conceptualization: MT, AT, KU

Methodology: MT, AT, KU

Investigation: MT, AT, LD, HC, MH, CT

Visualization: MT

Funding acquisition: KU, MT

Project administration: KU

Supervision: KU

Writing – original draft: MT

Writing – review & editing: All Authors

## Acknowledgements

This material is partially based upon work supported by the National Science Foundation under Grant No. EEC 1941543. Support from the US National Institutes of Health is gratefully acknowledged for the following awards: R01DK114506, R01DK096075, R01EB028782.

## Additional Information

Some authors declare competing interests. Drs. Uygun, Tessier, Yeh and Toner have patent applications relevant to this study. Drs. Uygun, Tessier and Toner have a financial interest in and serve on the Scientific Advisory Board for Sylvatica Biotech Inc., a company focused on developing high subzero organ preservation technology. Competing interests for MGH investigators are managed by the MGH and MGB in accordance with their conflict-of-interest policies.

---

**Supplementary Figure 1:**
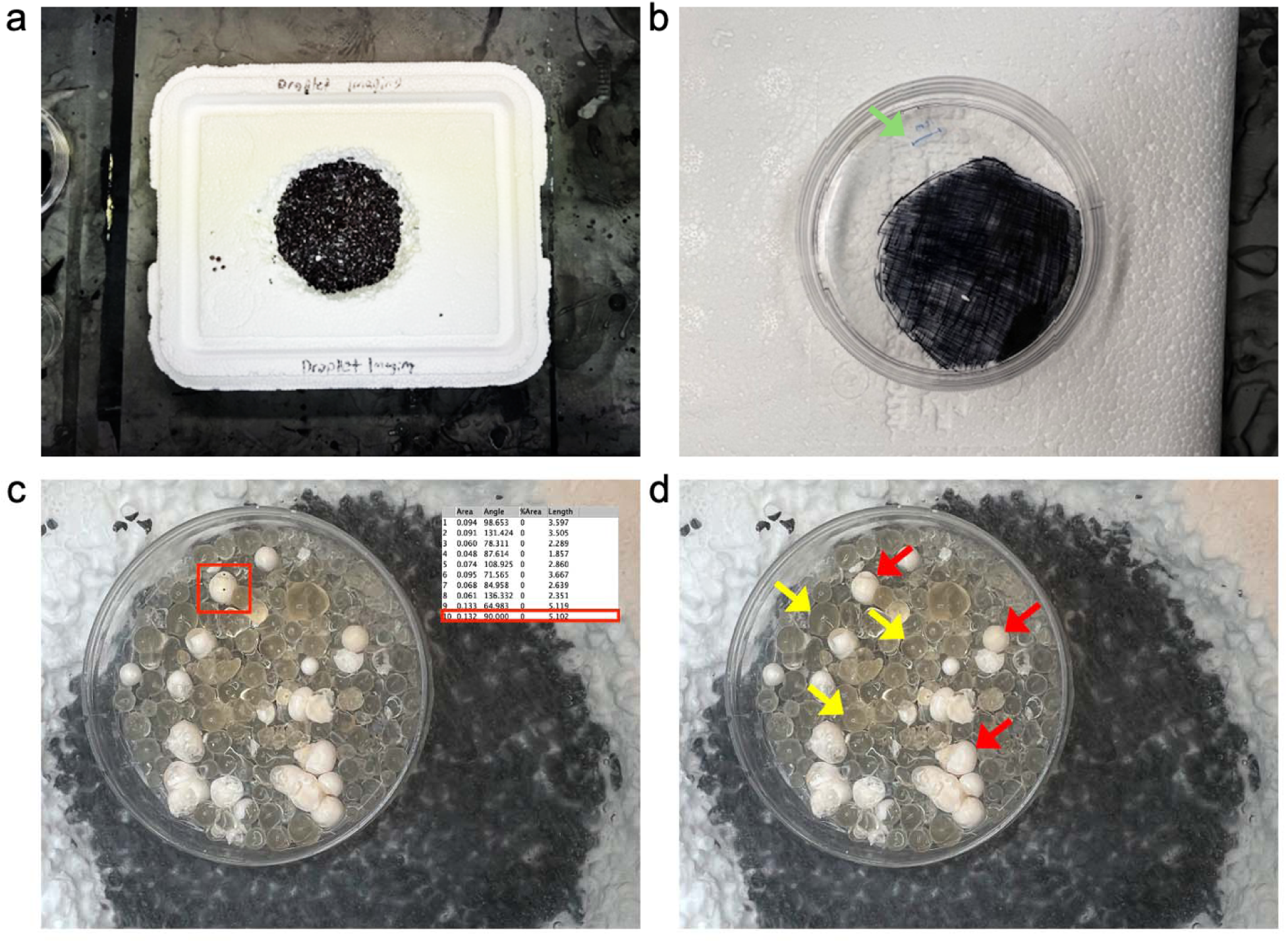
Characterization of Vitrified Droplets. To image the droplets while keeping them vitrified, a cryogenic environment had to be created. A divot capable of holding liquid nitrogen was cut out of a Styrofoam box and blackened to allow for contrast between the droplets and Styrofoam **(a)**. Into this divot was placed a blackened petri dish containing an etched scale, highlighted by the green arrow **(b)**. The petri dish was suspended on top of the liquid nitrogen and a thin film of LN_2_ was added inside, after which the droplets were poured in and imaged. The images were processed using FIJI, and the diameter of each droplet was manually determined; an example droplet size is outlined in red **(c)**. Each droplet was also marked based on whether it was vitrified or frozen, to allow for a direct size comparison; yellow arrows are vitrified droplets, while red arrows are frozen **(d)**.

**Supplementary Figure 2:**
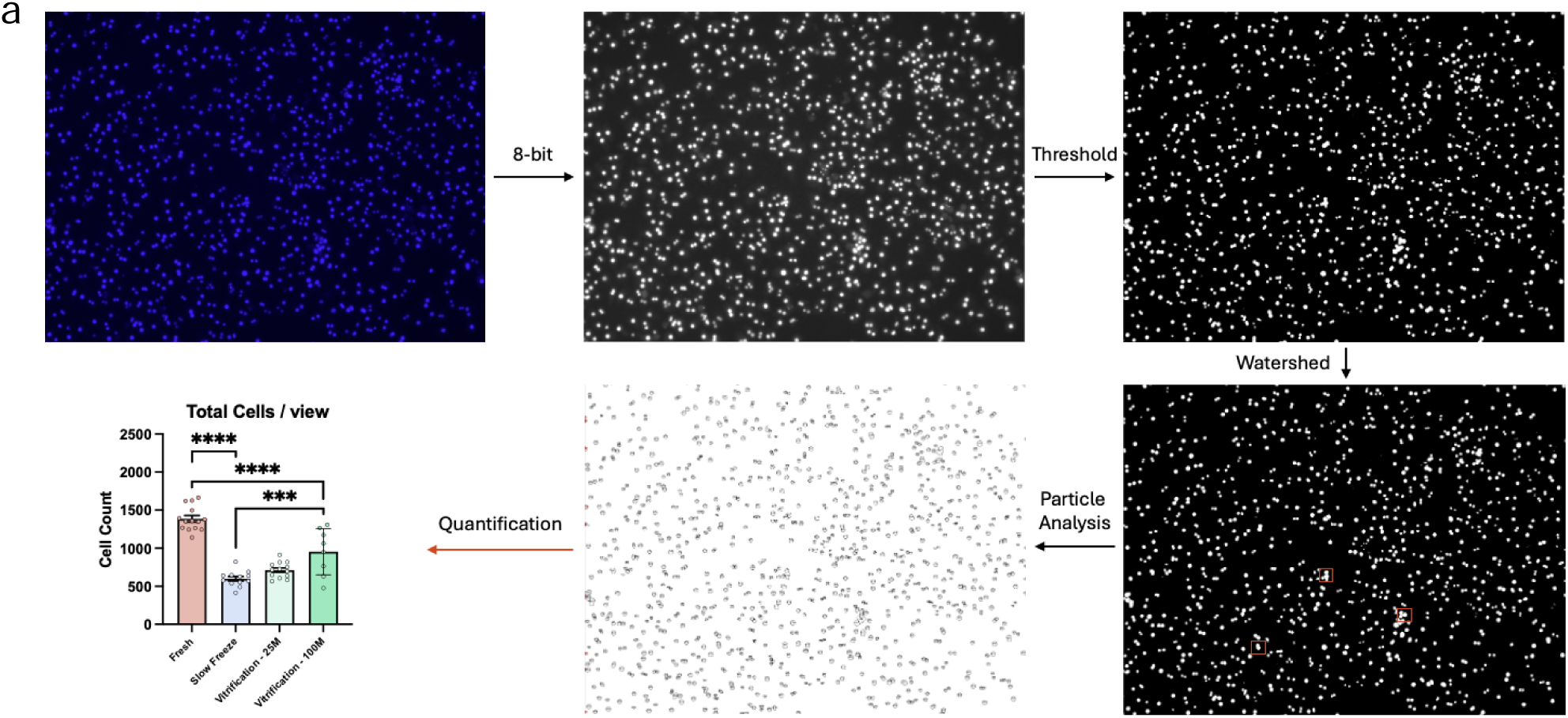
Image Analysis Workflow. Cells stained with NucBlue and Ethidium homodimer-1 were quantified to determine the total number of cells stained by each **(a)**. Color images were converted to 8-bit, after which a threshold was manually determined and applied to each image to eliminate false-positives and background noise. The watershed function was then implemented to segment any cells that overlapped; example segmentation is shown in red. The processed images then underwent particle analysis with a size range of 10-infinite pixels. This process was run on 2 images for both stains in each well as technical replicates, for all 4 groups.

**Supplementary Figure 3:**
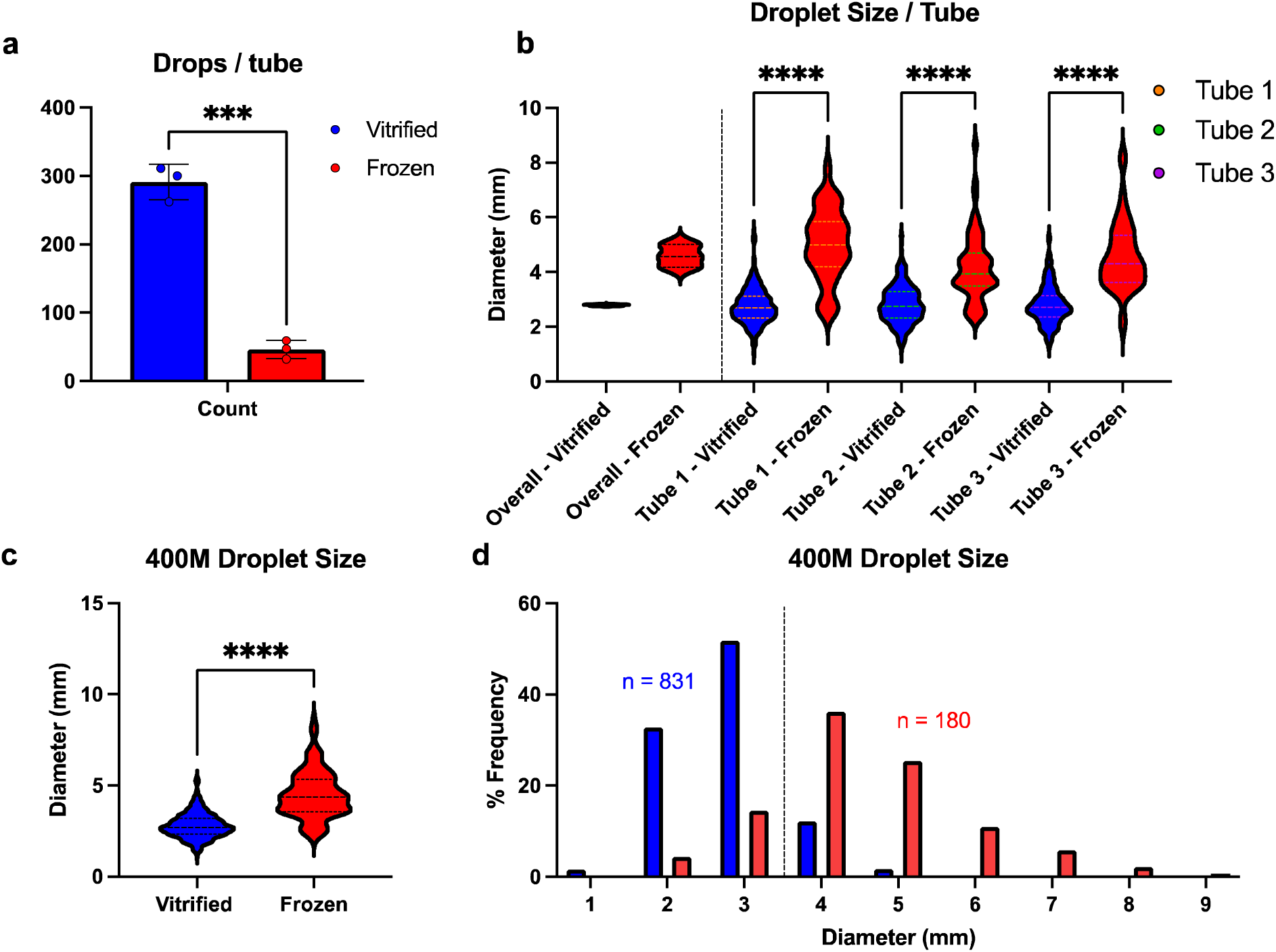
Droplet Characterization for 400M cells. Characterization of droplets obtained through parallelized droplet vitrification of the whole rat liver. Tubes showed an average vitrified count of 337 ± 39.0 droplets, constituting 86.6% ± 2.5 of the sample population (n = 4) **(a)**. Droplet size showed a consistent trend across all four tubes, with an average size difference of 64% ± 17 between vitrified and frozen droplets **(b)**. When comparing vitrified droplets (2.8mm ± 0.03) to frozen droplets (4.6 ± 0.4), a significant size difference is observed (p < 0.0001) **(c)**. A distinct size threshold of 4mm for vitrified droplets persists **(d)**. Results are displayed as mean ± standard deviation.

